# The FACTS model of speech motor control: fusing state estimation and task-based control

**DOI:** 10.1101/543728

**Authors:** Benjamin Parrell, Vikram Ramanarayanan, Srikantan Nagarajan, John Houde

## Abstract

We present a new computational model of speech motor control: the Feedback-Aware Control of Tasks in Speech or *FACTS* model. This model is based on a state feedback control architecture, which is widely accepted in non-speech motor domains. The FACTS model employs a hierarchical observer-based architecture, with a distinct higher-level controller of speech tasks and a lower-level controller of speech articulators. The task controller is modeled as a dynamical system governing the creation of desired constrictions in the vocal tract, based on the Task Dynamics model. Critically, both the task and articulatory controllers rely on an internal estimate of the current state of the vocal tract to generate motor commands. This internal state estimate is derived from initial predictions based on efference copy of applied controls. The resulting state estimate is then used to generate predictions of expected auditory and somatosensory feedback, and a comparison between predicted feedback and actual feedback is used to update the internal state prediction. We show that the FACTS model is able to qualitatively replicate many characteristics of the human speech system: the model is robust to noise in both the sensory and motor pathways, is relatively unaffected by a loss of auditory feedback but is more significantly impacted by the loss of somatosensory feedback, and responds appropriately to externally-imposed alterations of auditory and somatosensory feedback. The model also replicates previously hypothesized trade-offs between reliance on auditory and somatosensory feedback in speech motor control and shows for the first time how this relationship may be mediated by acuity in each sensory domain. These results have important implications for our understanding of the speech motor control system in humans.

## Introduction

Producing speech is one of the most complex motor activities humans perform. To produce even a single word, the activity of over 100 muscles must be precisely coordinated in space and time. This precise spatiotemporal control is difficult to master, and is not fully adult-like until the late teenage years^1^. How the brain and central nervous system (CNS) controls this complex system remains an outstanding question in speech motor neuroscience.

Early models of speech relied on servo control^2^. In this type of *feedback control* schema, the current feedback from the *plant* (thing to be controlled–for speech, this would be the articulators of the vocal tract, as well as perhaps the phonatory and respiratory systems) is compared against desired feedback and any discrepancy between the current and desired feedback drives the generation of motor commands to move the plant towards the current production goal. A challenge for any feedback control model of speech is the short, rapid movements that characterize speech motor behavior, with durations in the range of 50-300 ms. This is potentially shorter than the delays in the sensory systems. For speech, measured latencies to respond to external perturbations of the system range from 20-50 ms for unexpected mechanical loads^3, 4^ to around 150 ms for auditory perturbations^5, 6^. Therefore, the information about the state of the vocal tract conveyed by sensory feedback to the CNS is delayed in time. Such delays can cause huge problems for feedback control, leading to unstable movements and oscillations around goal states. Furthermore, speech production is possible even in the absence of auditory feedback, as seen in the ability of healthy speakers to produce speech when auditory and feedback is masked by loud noise^7, 8^. All of the above factors strongly indicate that speech cannot be controlled purely based on feedback control.

Several alternative approaches have been suggested to address these problems with feedback control in speech production and other motor domains. One approach, the equilibrium point hypothesis^9–11^, relegates feedback control to short-latency spinal or brainstem circuits operating on proprioceptive feedback, with high-level control based on pre-planned feedforward motor commands. Speech models of this type, such as the GEPPETO model, are able to reproduce many biomechanical aspects of speech but are not sensitive to auditory feedback^12–16^. Another approach is to combine feedback and feedforward controllers operating in parallel17, 18. This is the approach taken by the DIVA model^19–22^, which combines feedforward control based on desired articulatory positions with auditory and somatosensory feedback controllers. In this way, DIVA is sensitive to sensory feedback (via the feedback controllers) but capable of producing fast movements despite delayed or absent sensory feedback (via the feedforward controller).

A third approach, widely used in motor control models outside of speech, relies on the concept of state feedback control^23–26^. In this approach, the plant is assumed to have a state that is sufficiently detailed to predict the future behavior of the plant, and a controller drives the state of the plant towards a goal state, thereby accomplishing a desired behavior. A key concept in state feedback control is that the true state of the plant is not known to the controller; instead, it is only possible to estimate this state from efference copy of applied controls and sensory feedback. The internal state estimate is computed by first predicting the next plant state based on the applied controls. This state prediction is then used to generate predictions of expected feedback from the plant, and a comparison between predicted feedback and actual feedback is then used to correct the state prediction. Thus, in this process, the actual feedback from the plant only plays an indirect role in that it is only one of the inputs used to estimate the current state, making the system robust to feedback delays and noise.

We have earlier proposed a speech-specific instantiation of a state feedback control system^27^. The primary purpose of this earlier work was to establish the plausibility of the state feedback control architecture for speech production and suggest how such an architecture may be implemented in the human central nervous system. Computationally, our previous work built on models that have been developed in non-speech motor domains^24, 25^. Following this work, we implemented the state estimation process as a prototypical Kalman filter^28^, which provides an optimal posterior estimate of the plant state given a prior (the efference-copy-based state prediction) and a set of observations (the sensory reafference), assuming certain conditions such as a linear system. We subsequently implemented a one-dimensional model of vocal pitch control based on this framework^29^.

However, the speech production system is substantially more complex than our one-dimensional model of pitch. First, speech production requires the multi-dimensional control of redundant and interacting articulators (e.g., lips, tongue tip, tongue body, jaw, etc.). Second, speech production relies on the control of high-level task goals rather than direct control of the articulatory configuration of the plant (e.g., for speech, positions of the vocal tract articulators). For example, speakers are able to compensate immediately for a bite block which fixes the jaw in place, producing essentially normal vowels^30^. Additionally, speakers react to displacement of a speech articulator by making compensatory movements of other articulators: speakers lower the upper lip when the jaw is pulled downward during production of a bilabial [b]^3, 4^, and raise the lower lip when the upper lip is displaced upwards during production of [p]^31^. Importantly, these actions are not reflexes, but are specific to the ongoing speech task. No upper lip movement is seen when the jaw is displaced during production of [z] (which does not require the lips to be close), nor is the lower lip movement increased if the upper lip is raised during production of [f] (where the upper lip is not involved). Together, these results strongly indicate that the goal of speech is not to achieve desired positions of each individual speech articulator, but must rather be to achieve some higher-level goal. While most models of speech motor production thus implement control at a higher speech-relevant level, the precise nature of these goals (vocal tract constrictions^32–34^, auditory patterns^2, 12, 21^, or both^14, 22^) remains an ongoing debate.

One prominent model that employs control of high-level speech tasks rather than direct control of articulatory variables is the Task Dynamic model^32, 35^. In Task Dynamics, the state of the plant (current positions and velocities of the speech articulators) is assumed to be available through proprioception. Importantly, this information is not used to directly generate an error or motor command. Rather, the current state of the plant is used to calculate values for various constrictions in the vocal tract (e.g., the distance between the upper and lower lip, the distance between the tongue tip and palate, etc.). It is these *constrictions*, rather than the positions of the individual articulators, that constitute the goals of speech production in Task Dynamics.

The model proposed here (Feedback Aware Control of Tasks in Speech, or *FACTS*) extends the idea of articulatory state estimation from the simple linear pitch control mechanism of our previous SFC model to the highly non-linear speech articulatory system. This presents three primary challenges: first, moving from pitch control to articulatory control requires the implementation of control at a higher level of speech-relevant tasks, rather than at the simpler level of articulator positions. To address this issue, FACTS is built upon the Task Dynamics model, as described above. However, unlike the Task Dynamics model, which assumes the state of the vocal tract is directly available through proprioception, here we model the more realistic situation in which the vocal tract state must be estimated from an efference copy of applied motor commands as well as somatosensory and auditory feedback. The second challenge is that this estimation process is highly non-linear. This required that the implementation of the observer as a Kalman filter in SFC be altered, as this estimation process is only applicable to linear systems. Here, we implement state estimation as an Unscented Kalman Filter^36^, which is able to account for the nonlinearities in the speech production system and incorporates internal state prediction, auditory feedback, and somatosensory feedback. Lastly, the highly non-linear mapping between articulatory positions and speech acoustics must be approximated in order to predict the auditory feedback during speech. Here, we learn the articulatory-to-acoustic mapping using Locally Weighted Projection Regression or LWPR^37^. Thus, in the proposed FACTS model, we combine the hierarchical architecture and task-based control of Task Dynamics with a non-linear state-estimation procedure to develop a model that is capable of rapid movements with or without sensory feedback, yet is still sensitive to external perturbations of both auditory and somatosensory feedback.

In the following sections, we describe the architecture of the model and the computational implementation of each model process. We then describe the results of a number of model simulations designed to test the ability of the model to simulate human speech behavior. First, we probe the behavior of the model when sensory feedback is limited to a single modality (somatosensation or audition), removed entirely, or corrupted by varying amounts of noise. Second, we test the ability of the model to respond to external auditory or mechanical perturbations. In both of these sections, we show that the model performs similarly to human speech behavior and makes new, testable predictions about the effects of sensory deprivation. Lastly, we explore how sensory acuity in both the auditory and somatosensory pathways affects the model’s response to external auditory perturbations. Here, we show that the response to auditory perturbations depends not only on auditory acuity but on somatosensory acuity as well, demonstrating for the first time a potential computational principle that may underlie the demonstrated trade-off in response magnitude to auditory and somatosensory perturbations across human speakers.

### Model Overview

A schematic control diagram of the FACTS model is shown in Fig. 1. Modules that build on Task Dynamics are shown in blue, and the articulatory state estimation process (or observer) is shown in red. Following Task Dynamics, speech tasks in the FACTS model are hypothesized to be desired constrictions in the vocal tract (e.g., close the lips for a [b]). Each of these speech tasks, or gestures, can be specified in terms of it’s constriction location (where in the vocal tract the constriction is formed) and it’s constriction degree (how narrow the constriction is). We model each gesture as a separate critically-damped second-order system^32^. Interestingly, similar dynamical behavior has been seen at a neural population level during the planning and execution of reaching movements in non-human primates^39, 40^, suggesting that a dynamical systems model of task-level control may be an appropriate first approximation to the neural activity that controls movement production. However, the architecture of the model would also allow for tasks in other control spaces, such as auditory targets (c.f.^13, 19^), though an appropriate task feedback control law^1^ for such targets would need to be developed.

**Figure 1.**
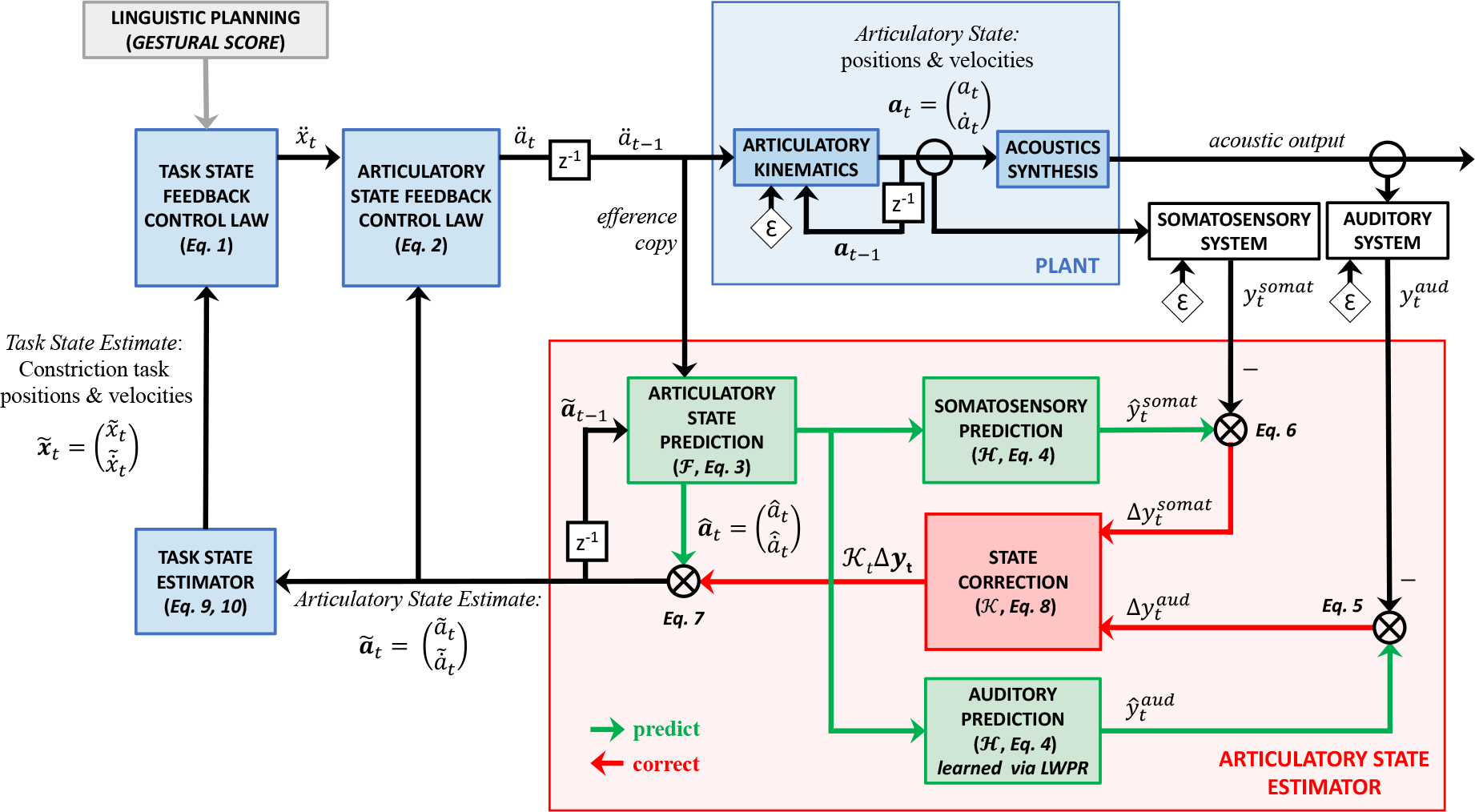
Architecture of the FACTS model. Boxes in blue represent processes outlined in the Task Dynamics model^32, 38^. The articulatory state estimator (shown in red) is implemented as an Unscented Kalman Filter, which estimates the current articulatory state of the plant by combining the predicted state generated by a forward model with auditory and somatosensory feedback. Additive noise is represented by *ɛ*. Time-step delays are represented by *z*^−1^. Equation numbers correspond to equations found in the methods.

FACTS uses as the Haskins Configurable Articulatory Synthesizer (or CASY)^38, 41, 42^ as the model of the vocal tract **plant** being controlled. The relevant parameters of the CASY model required to move the tongue body to produce a vowel (and which fully describe the articulatory space for the majority of the simulations in this paper) are the Jaw Angle (JA, angle of the jaw relative to the temporomandibular joint), Condyle Angle (CA, the angle of the center of the tongue relative to the jaw, measured at the temporomandibular joint), and the Condyle Length (CL, distance of the center of the tongue from the temporomandibular joint along the Condyle Angle). The CASY model is shown in Fig. 2.

**Figure 2.**
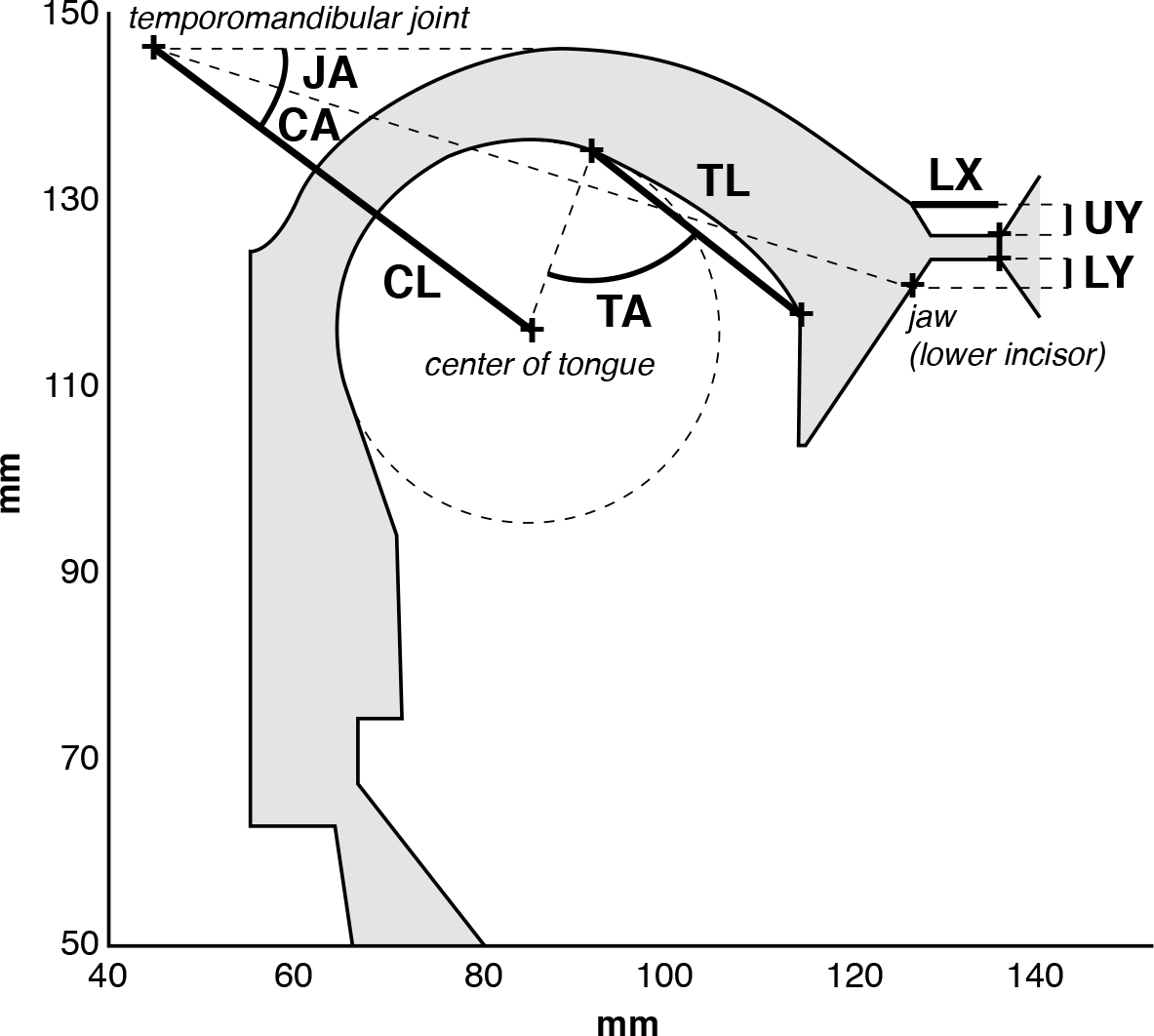
The CASY plant model. Articulatory variable relevant to the current paper are the Jaw Angle (JA), Condyle Angle (CA), and Condyle Length (CL). See text for a description of these variables. Diagram after^43^.

The model begins by receiving the output from a linguistic planning module. Currently, this is implemented as a *gestural score* in the framework of Articulatory Phonology^33, 34^. These gestural scores list the control parameters (e.g., target constriction degree, constriction location, damping, etc.) for each gesture in a desired utterance as well as each gesture’s onset and offset times. For example, the word “mod" ([mAd]) has four gestures: simultaneous activation of a gesture driving closure at the lips for [m], a gesture driving an opening of the velum for nasalization of [m], and a gesture driving a wide opening between the tongue and hard palate for the vowel [A]. These are followed by a gesture driving closure between the tongue tip and hard palate for [d] (Fig. 3).

**Figure 3.**
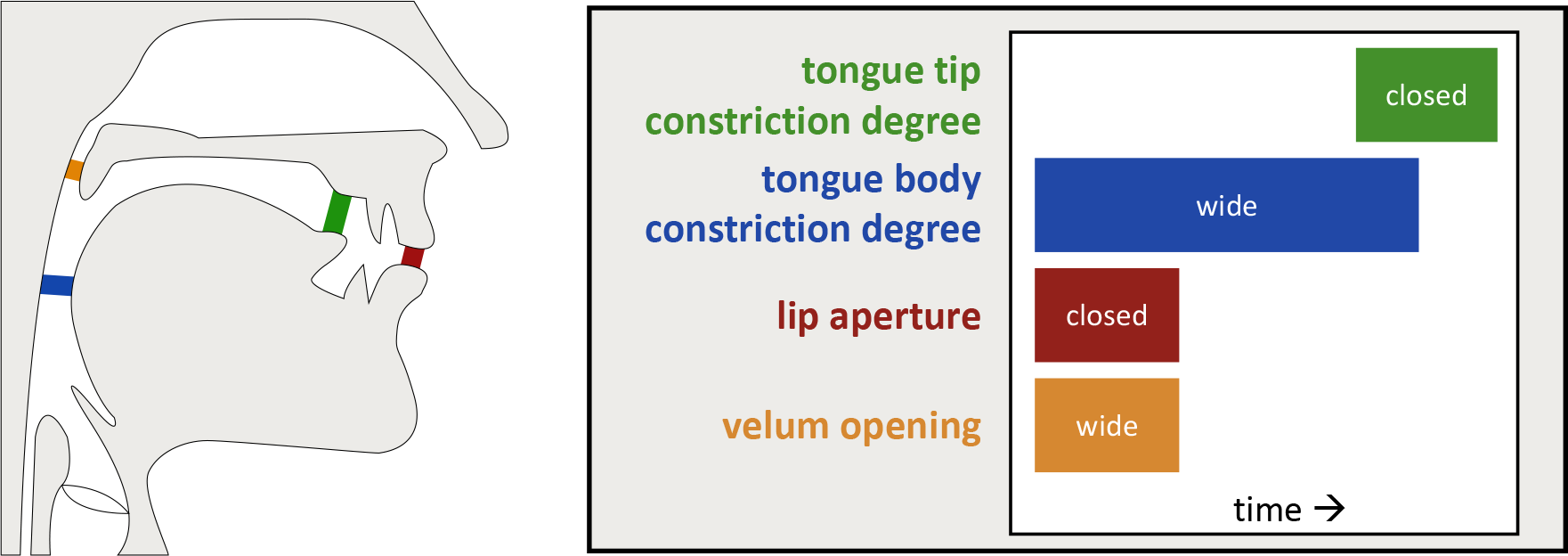
Example of task variables and gestural score for the word “mod”. A gestural score for the word “mod” [mAd], which consists of a bilabial closure and velum opening gestures for [m], a wide constriction in the pharynx for [A], and a tongue tip closure gesture for [d]. Tasks are shown on the left, and a schematic of the gestural score on the right.

The **task state feedback control law** takes these gestural scores as input and generates a task-level command based on the current state of the ongoing articulatory tasks. In this way, the task-level commands are dependent on the current task-level state. For example, if the lips are already closed during production of a /b/, a very different command needs to be generated than if the lips are far apart. These task-level commands are converted into motor commands that can drive changes in the positions of the speech articulators by the **articulatory state feedback control law**, using information about the current articulatory state of the vocal tract. The motor commands generate changes in the model vocal tract articulators (or **plant**), which are then used to generate an acoustic signal.

The **articulatory state estimator**(sometimes called an observer in other models) combines a copy of the outgoing motor command (or efference copy) with auditory and somatosensory feedback to generate an internal estimate of the articulatory state of the plant. First, the efference copy of the motor command is used (in combination with the previous arituclatory state estimate) to generate a prediction of the articulatory state. This is then used by a **forward model** (learned via LWPR) to generate auditory and somatosensory predictions, which are compared to incoming sensory signals to generate sensory errors. Subsequently, these sensory errors are used to correct the state prediction to generate the final state estimate.

The final articulatory state estimate is used by the articulatory state feedback control law to generate the next motor command, as well as being passed to the **task state estimator** to estimate the current task state, or values (positions) and first derivatives (velocities) of the speech tasks (note the Task State was called the Vocal Tract State in earlier presentations of the model^44, 45^). Finally, this estimated task-level state is passed to the task state feedback control law to generate the next task-level command.

A more detailed mathematical description of the model can be found in the methods.

## Results

Here we present results showing the accuracy of the learned forward model, and of various modeling experiments designed to test the ability of the model to qualitatively replicate human speech motor behavior under various conditions, including both normal speech as well as externally perturbed speech.

### Forward model accuracy

Figure 4 visualizes a three dimensional subspace of the learned mapping from the 10-dimensional articulatory state space to the 3-dimensional space of formant frequencies (F1 – F3). Specifically, we look at the mapping from the tongue condyle length and condyle angle to the first (see Figure 4A-C) and second formants (see Figure 4D-F), projected onto each two-dimensional plane. We also plot normalized histograms of the number of receptive fields that cover each region of the space (represented as a heatmap in Figure 4A and D and with a thick blue line in the other subplots). In each figure, the size of the circles is proportional to the absolute value of the error between the actual and predicted formant values. Overall, the fit of the model is very good, with an average error of 4.2 Hz (std., 9.1 Hz) for F1 and 6.6 Hz (std., 19.1 Hz) for F2. Fit error increases in regions of the space that are relatively sparsely covered by receptive fields. In addition, the higher frequency of smaller circles at the margins of the distribution (and therefore the edges of the articulatory space) suggest that we need fewer receptive fields to cover these regions. Of course, this means that we do see some bigger circles in these regions where the functional mapping is not adequately represented by a small number of fields. Also note that we are only plotting the number of receptive fields that are employed to cover a given region of articulatory space, and this is *not* indicative of how much weight they carry in representing that region of space.

**Figure 4.**
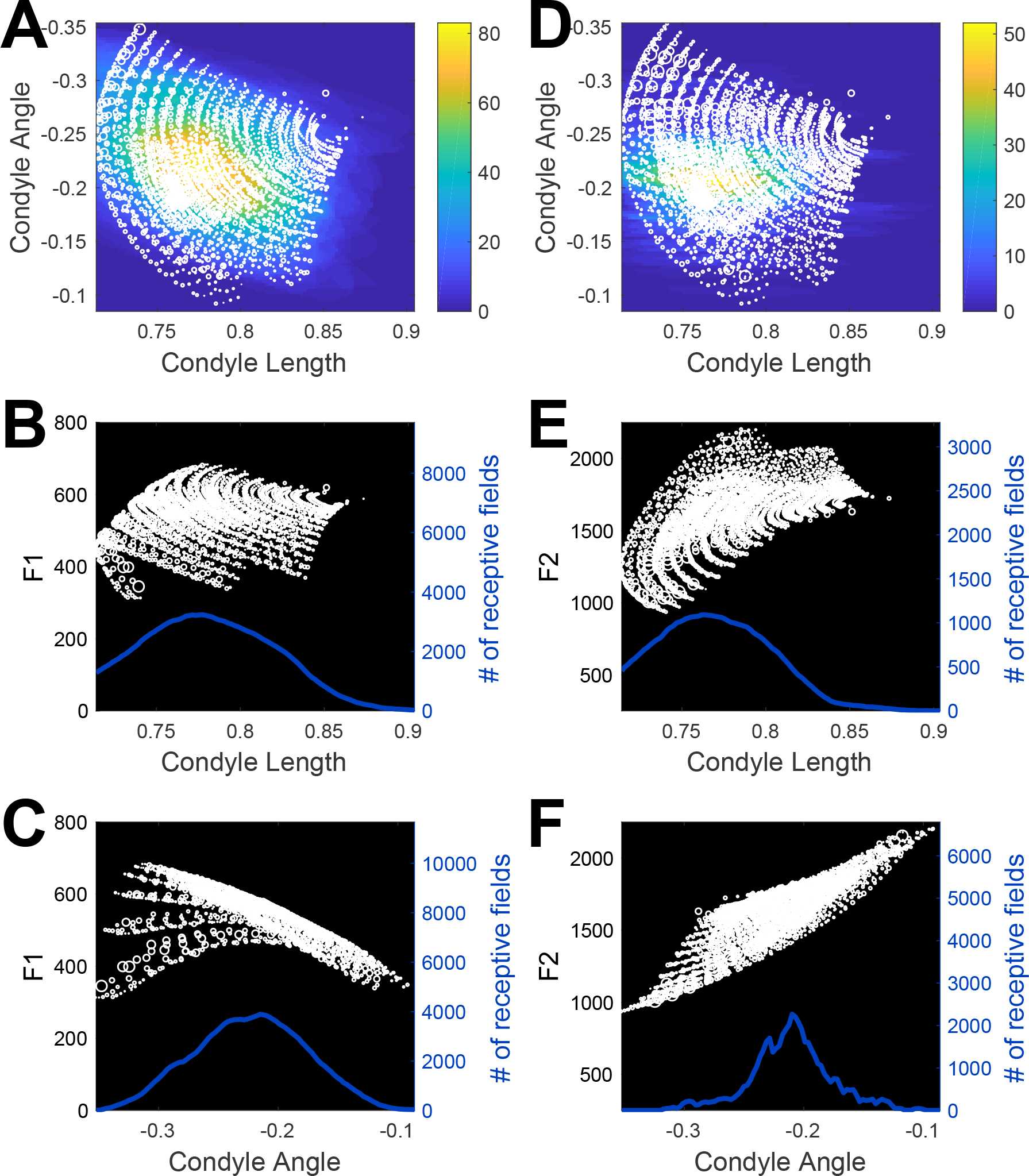
Learned LWPR transformations from CASY articulatory parameters to acoustics. Predicted formant values are shown as white circles. The width of each circle is proportional to the absolute value of the error between the actual formant values and the formant values predicted by the LWPR model. The density distributions reflect the number of receptive fields that cover each point (represented as colors in A, D and the height of the line in B, C, D, F). (A-C) show F1 values. (D-F) show F2 values. (A) Condyle Angle vs Condyle Length, (B) F1 vs Condyle Length, (C) F1 vs Condyle Angle. (D-F), replicate (A-C) for F2.

### Model response to changes in sensory feedback availability and noise levels

How does the presence or absence of sensory feedback affect the speech motor control system? While there is no direct evidence to date on the effects of total loss of sensory information in human speech, some evidence comes from when sensory feedback from a single modality is attenuated or eliminated. Notably, the effects of removing auditory and somatosensory feedback differ. In terms of auditory feedback, speech production is relatively unaffected by it’s absence: speech is generally unaffected by when auditory feedback is masked by loud masking noise^7, 8^. However, alterations to somatosensory feedback have larger effects: blocking oral tactile sensation through afferent nerve injections or oral anesthesia leads to substantial imprecision in speech articulation^46, 47^.

Fig. 5 presents simulations from the FACTS model testing the ability of the model to replicate the effects of removing sensory feedback seen in human speech. All simulations modeled the vowel sequence [ǝ a i]. 100 simulations were run for each of four conditions: normal feedback (5B), somatosensory feedback only (5C), auditory feedback only (5D), and no sensory feedback (5E). For clarity, only the trajectory of the tongue body in the CASY articulatory model is shown for each simulation.

**Figure 5.**
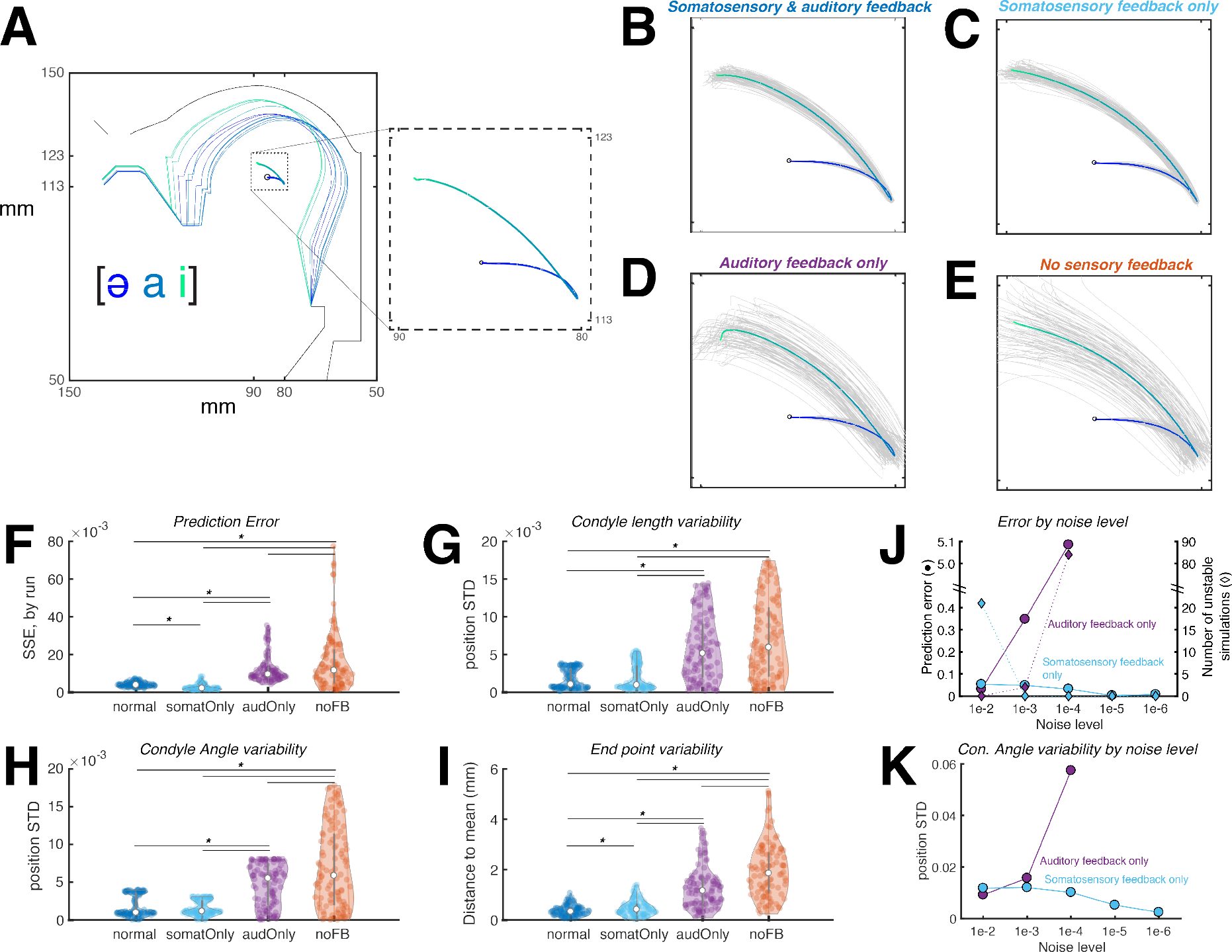
FACTS simulation of the vowel sequence [ǝ a i], varying the type of sensory feedback available to the model. (A (shows a sample simulation with movements of the CASY model as well as the trajectory of the tongue center. (B-E) each show tongue center trajectories from 100 simulations (gray) and their mean (blue-green gradient) with varying types of sensory feedback available. Variability is lower when sensory feedback is available, and lower when auditory feedback is absent compared to when somatosensory feedback is absent. (F) shows the prediction error in each condition. (G-H) show the produced variability in two articulatory parameters of the CASY plant model related to vowel production and (I) shows variability of the tongue center at the final simulation sample. (J) and (K) show prediction error and articulatory variability relative to sensory noise levels when only one feedback channel is available. Decreasing sensory noise leads to increased accuracy for somatosensation but decreased accuracy for audition. Colors in (F-K) correspond to the colors in the titles of (B-E).

In the normal feedback condition (Fig. 5B), the tongue lowers from [ǝ] to [a], then raises and fronts from [a] to [i]. Note that there is some variability across simulation runs due to the noise in the motor and sensory systems. This variability is also found in human behavior and the ability of the state feedback control architecture to replicate this variability is a strength of this approach25.

The effect of removing auditory feedback (Fig. 5C) leads to a significant, though small, increase in the variability of the tongue body movement as measured by the tongue location at the movement endpoint (see Fig. 5I), though this effect was not seen in measures of Condyle Angle or Condyle Length variability (5G-H). Interestingly, while variablity increased, prediction error slightly decreased in this condition 5F. Overall, these results are consistent with experimental results that demonstrate that speech is essentially unaffected, in the short term, by the loss of auditory information (though auditory feedback is important for pitch and amplitude regulation^48^ as well as to maintain articulatory accuracy in the long term^48–50^).

Removing somatosensory feedback while maintaining auditory feedback (Fig. 5B) leads to an increase in both variability across simulation runs as well as an increase in prediction error (Fig. 5F-I). This result is broadly consistent with the fact that reduction of tactile sensation via oral anaesthetic or nerve block leads to imprecise articulation for both consonants and vowels^47, 51^ (though the acoustic effects of this imprecision may be less perceptible for vowels^47^). However, a caveat must be made that our current model does not include tactile sensation, only proprioceptive information. Unfortunately, it is impossible to block proprioceptive information from the tongue, as that afferent information is likely carried along the same nerve (hypoglossal nerve) as the efferent motor commands^52^. It is difficult to prove, then, exactly how a complete loss of proprioception would affect speech. Nonetheless, the current model results are consistent with studies that how shown severe dyskinesia in reaching movements after elimination of proprioception in non-human primates (see^53^ for a review) and in human patients with severe proprioceptive deficits^54^. In summary, although the FACTS model currently includes only proprioceptive sensory information rather than both proprioceptive and tactile signals, these simulation results are consistent with a critical role for the somatosensory system in maintaining the fine accuracy of the speech motor control system.

While removal of only auditory feedback lead to only small increases in variability (in both FACTS simulations and human speech), our simulations show speech in the complete absence of sensory feedback (Fig. 5E) shows much larger variability than the absence of either auditory or somatosensory feedback alone. This is consistent with human behvaior^51^, and occurs because without sensory feedback there is no way to detect and correct for the noise inherent in the motor system (shown by the large prediction errors and increased articulatory variability in Fig. 5F.

The effects of changing the noise levels in the system can be see in Fig. 5J-K. For these simulations, only one type of feedback was used at a time: somatosensory (cyan) or auditory (purple). Noise levels (shown on the x axis) reflect both the sensory system noise and the internal estimate of that noise, which were set to be equal. Each data point reflects 100 stable simulations. Data for the acoustic-only simulations are not shown for noise levels below 1e-5 as the model became highly unstable in these conditions to due inaccurate articulatory state estimates (the number of unstable or divergent simulations is shown in Fig. 5J). For the somatosensory system, the prediction error and articulatory variability (shown here for the Condyle Angle) *decrease* as the noise decreases. However, for the auditory system, both prediction error and articulatory variability *increase* as the noise decreases. Because of the Kalman gain, decreased noise in a sensory or predictive signal leads not only to a more accurate signal, but also to a greater reliance on that signal compared to the internal state prediction. When the system relies more on the somatosensory signal, this results in a more accurate state estimate as the somatosensory signal directly reflects the state of the plant. When the system relies more on the auditory signal, however, this results in an less accurate state estimate as the auditory signal only indirectly reflects the state of the plant as a results of the nonlinear, many-to-one articulatory-to-acoustic mapping of the vocal tract.

In sum, FACTS is able to replicate the variability seen in human speech, as well as qualitatively match the effects of both auditory and somatosensory masking on speech accuracy. While the variability of human speech in the absence of proprioceptive feedback remains untested, the FACTS simulation results make a strong prediction that could be empirically tested in future work if some manner of blocking or altering proprioceptive signals could be devised.

### Model response to mechanical and auditory perturbations

When a downward mechanical load is applied to the jaw during the production of a consonant, speakers respond by increasing the movements of the other speech articulators in a task-specific manner to achieve full closure of the vocal tract^3, 4, 31^. For example, when the jaw is perturbed during production of a bilabial stops /b/ or /p/, the upper lip moves downward to a greater extent than normal to compensate for the lower jaw position. This upper lip lowering is not found for jaw perturbations during /f/ or /z/, indicating it is specific to sounds produced using the upper lip. Conversely, tongue muscle activity is larger following jaw perturbation for /z/, which involves a constriction made with the tongue tip, but not for /b/, for which the tongue is not actively involved.

The ability to sense and compensate for mechanical perturbations relies on the somatosensory system. We tested the ability of FACTS to reproduce the task-specific compensatory responses to jaw load application seen in human speakers by applying a small downward acceleration to jaw (Jaw Angle parameter in CASY) starting midway through a consonant closure for stops produced with the lips (/p/) and tongue tip (/t/). The perturbation continued to the end of the word. As shown in Fig. 6A, the model produces greater lowering of the upper lip (as well as greater raising of the lower lip) when the jaw is fixed during production of /p/, but not during /t/, mirroring the observed response in human speech.

**Figure 6.**
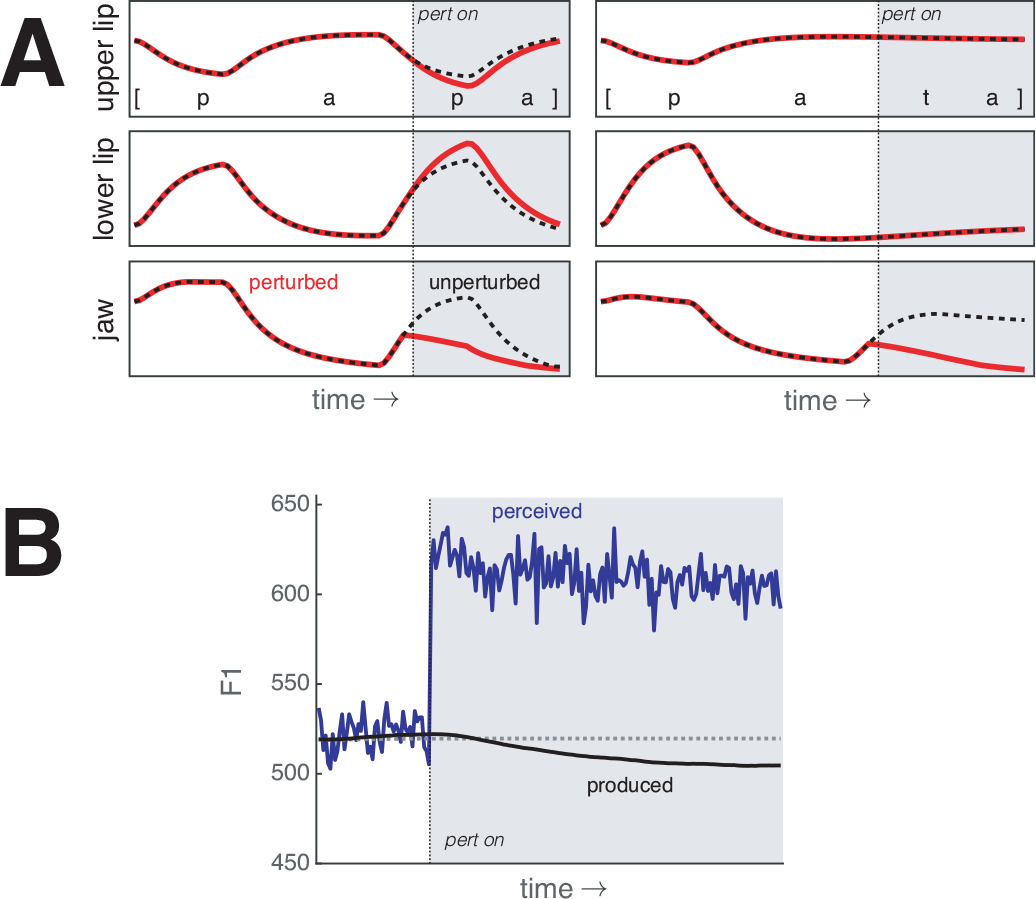
FACTS model simulations of mechanical and auditory perturbations. Times when the perturbations are applied are shown in gray. (A) shows the response of the model to fixing the jaw in place (simulating a downward force applied to the jaw) midway through the closure for second consonant in [papa] (left) and [pata] (right). Unperturbed trajectories are indicated with a dashed black line and perturbed trajectories with a solid red line. The upper and lower lips move more to compensate for the jaw perturbation only when the perturbation occurs on [p], mirroring the task-specific response seen in human speakers. (B) shows the response of the FACTS model to a +100 Hz auditory perturbation of F1 while producing [ǝ]. The produced F1 is shown in black and the perceived F1 is shown in blue. Note that the perceived F1 is corrupted by noise. The model responds to the perturbation by lowering F1 despite the lack of an explicit auditory target. The partial compensation to the perturbation produced by the model matches that observed in human speech.

In addition to the task-specific response to mechanical perturbations, speakers will also adjust their speech in response to auditory perturbations^55^. For example, when the first vowel formant (F1) is artificially shifted upwards, speakers produce within-utterance compensatory responses by lowering their produced F1. The magnitude of these responses only partially compensates for the perturbation, unlike the complete responses produced for mechanical perturbations. While the exact reason for this partial compensation is not known, it has been hypothesized to relate to small feedback gains^19^ or conflict with the somatosensory feedback system^14^. We explore the cause of this partial compensation below, but focus here on the ability of the model to replicate the observed behavior.

To test the ability of the FACTS model to reproduce the observed partial responses to auditory feedback perturbations, we simulated production of a steady-state [ǝ] vowel. After a brief stabilization period, we abruptly introduced a +100 Hz perturbation of F1 by adding 100 Hz to the perceived F1 signal in the auditory processing stage. This introduced a discrepancy between the produced F1 (shown in black in Fig. 6B) and the perceived F1 (shown in blue in Fig. 6B). Upon introduction of the perturbation, the model starting to produce a compensatory lowering of F1, eventually reaching a steady value below the unperturbed production. This compensation, like the response in human speakers, is only partial (roughly 20 Hz or 15% of the total perturbation).

Importantly, FACTS produces compensation auditory perturbations despite having no auditory targets. Previously, such compensation has been seen as evidence in favor of the existence of auditory targets for speech^14^. In FACTS, auditory perturbations cause a change in the estimated state of the vocal tract on which the task-level and articulatory-level feedback controllers operate. This causes a change in motor behavior compared to the unperturbed condition, resulting in what appears to be compensation for the auditory perturbation. Our model results thus show that this compensation is possible without explicit auditory goals. Of course, these results do not argue that auditory goals do not exist. Rather, we show that they are not necessary for this behavior.

### Model trade-offs between auditory and somatosensory acuity

The amount of compensation to an auditory perturbation has been found to vary substantially between individuals^55^. One explanation for the inter-individual variability is that the degree of compensation is related to the acuity of the auditory system. Indeed, there seems to be a relationship between measurements of auditory acuity and the magnitude of the compensatory response to auditory perturbation of vowel formants^56^. If we assume that acuity is inversely related to the amount of noise in the sensory system, this explanation fits with the UKF implementation of the state estimation procedure in FACTS, where the weight assigned to the auditory error is ultimately related to the estimate of the noise in the auditory system. In Fig. 7B, we show that by varying the amount of noise in the auditory system (along with the internal estimate of that noise), we can drive differences in the amount of compensation the model produces to a +100 Hz perturbation of F1. When we double the auditory noise compared to baseline (top), the compensatory response is reduced. When we halve the auditory noise (bottom), the response increases.

**Figure 7.**
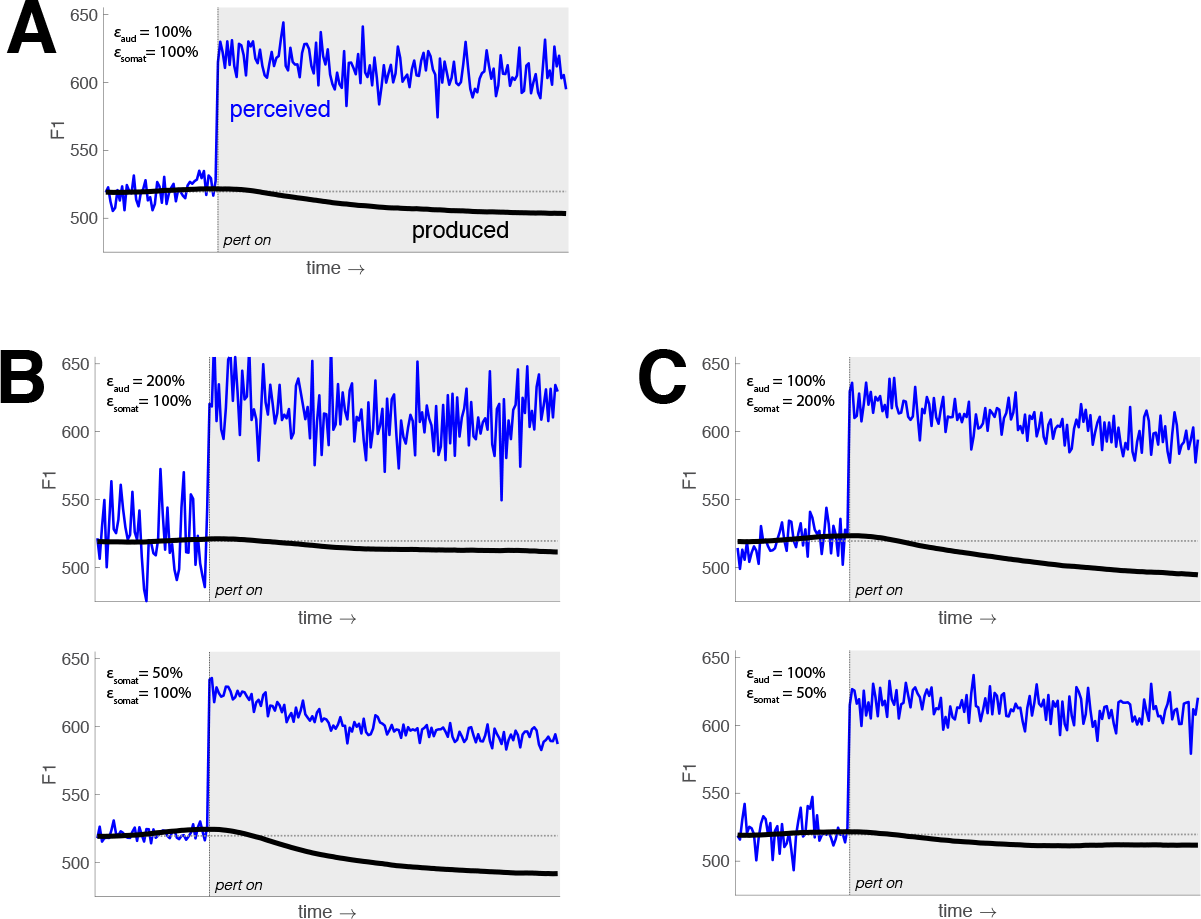
FACTS model simulations of the response to a +100 Hz perturbation of F1. (A) shows the baseline response. shows the effects of altering the amount of noise in the auditory system in tandem with the observer’s estimate of that noise. An increase in auditory noise (top) leads to a smaller perturbation response, while a decrease in auditory noise (bottom) leads to a larger response. (C) shows the effects of altering the amount of noise in the somatosensory system in tandem with the observer’s estimate of that noise. The pattern is opposite of that for auditory noise. Here, an increase in somatosensory noise (top) leads to a larger perturbation response, while a decrease in somatosensory noise (bottom) leads to a smaller response.

Interestingly, the math underlying the UKF suggests that the magnitude of the response to an auditory error should be tied not only to the acuity of the auditory system, but to the acuity of the somatosensory system as well. This is because the weights assigned by the Kalman filter take the full noise covariance of all sensory systems into account. We verified this prediction empirically by running a second set of simulated responses to the same +100 Hz perturbation of F1, this time maintaining the level of auditory noise constant while varying only the level of somatosensory noise. The results can be seen in Fig. 7C. When the level of somatosensory noise is increased, the response to the auditory perturbation increases. Conversely, when the level of somatosensory noise is reduced, the compensatory response is reduced as well. These results suggest that the compensatory response in human speakers should be related to the acuity of the somatosensory system as well as the auditory system, a novel hypothesis which we are currently testing experimentally. Broadly, however, these results agree with, and provide a testable hypothesis about the cause of, empirical findings that show a trading relationship across speakers in their response to auditory and somatosensory perturbations^57^.

## Discussion

The proposed FACTS model provides a novel way to understand the speech motor system. The model is an implementation of state feedback control that combines high-level control of speech tasks with a non-linear method for estimating the current articulatory state to drive speech motor behavior. We have shown that the model replicates many important characteristics of human speech motor behavior: the model produces stable articulatory behavior, but with some trial-to-trial variability. This variability increases when somatosensory information is unavailable, but is largely unaffected by the loss of auditory feedback.

The model is also able to reproduce task-specific responses to external perturbations. For somatosensory perturbations, when a downward force is applied to the jaw during production of an oral consonant, there is an immediate compensatory response *only in those articulators needed to produce the current task*. This is seen in the increased movement of the upper and lower lips to compensate for the jaw perturbation during production of a bilabial /b/ but no alterations in lip movements when the jaw was perturbed during production of a tongue-tip consonant /d/. The ability of the model to respond to perturbations in a task-specific replicates a critical aspect of human speech behavior and is due to the inclusion of a task state feedback control law in the model^35^.

For auditory perturbations, we showed that FACTS is able to produce compensatory responses to external perturbations of F1, even though there is no explicit auditory goal in the model. Rather, the auditory signal is used to inform the observer about the current state of the vocal tract articulators. We additionally showed that FACTS is able to produce the inter-individual variability in the magnitude of this compensatory response as well as the previously observed relationship between the magnitude of this response and auditory acuity.

We have also shown that FACTS makes some predictions about the speech motor system that go beyond what has been demonstrated experimentally to date. FACTS predicts that a complete loss of sensory feedback would lead to large increases in articulatory variability beyond those seen in the absence of auditory or somatosensory feedback alone. Additionally, FACTS predicts that the magnitude of compensation for auditory perturbations should be related not only to auditory acuity, but to somatosensory acuity as well. These concrete predictions can be experimentally tested to assess the validity of the FACTS model, testing which is ongoing in our labs.

One of the major drawbacks of the current implementation of FACTS is that the model of the plant only requires kinematic control of articulatory positions. While a kinematic approach is relatively widespread in the speech motor control field– including both DIVA and Task Dynamics–there is experimental evidence that the dynamic properties of the articulators, such as gravity and tissue elasticity, need to be accounted for in its control^58–61^. Moreover, speakers will learn to compensate for perturbations of jaw protrusion that are dependent on jaw velocity^57, 62–64^, indicating that speakers are able to generate motor commands that anticipate and cancel out the effects of those altered articulatory dynamics. While the FACTS model in its current implementation does not replicate this dynamical control of the speech articulators, the overall architecture of the model is compatible with control of dynamics rather than just kinematics^23^. Control of articulatory dynamics would require a dynamic model of the plant and implementing a new articulatory-level feedback control law that would output motor commands as forces, rather than (or potentially in addition to) articulatory accelerations. Coupled with parallel changes to the articulatory state prediction process, this would allow for FACTS to control a dynamical plant without any changes to the overall architecture of the model.

While a detailed discussion of the neural basis of the computations in FACTS is beyond the scope of the current paper, in order to demonstrate the plausibility of FACTS as a neural model of speech motor control, we briefly touch on potential neural substrates that may underlie a state-feedback control architecture in speech^23, 24, 27^. The cerebellum is widely considered to play a critical role as an internal forward model to predict future articulatory and sensory states^26, 65^. The process of state estimation may occur in the parietal cortex, and indeed inhibitory stimulation of the inferior parietal cortex with transcranial magnetic stimulation impairs sensorimotor learning in speech^66^, consistent with a role in this process. However, state estimation for speech may also (or alternatively) reside in the ventral premotor cortex (vPMC) for speech, where the premotor cortices are well situated for integrating sensory information (received from sensory corteces via the arcuate fasiculus and the superior longitudinal fasiculus) with motor efference copy from primary motor cortex and cerebellum. Primary motor cortex (M1), with its descending control of the vocal tract musculature and bidirectional monosynaptic connections to primary sensory cortex, is the likely location of the articulatory feedback control law, converting task-level commands from vPMC to articulatory motor commands. Interestingly, recent work using electrocorticography has shown that areas in M1 code activation of task-specific muscle synergies similar to those proposed in Task Dynamics and FACTS^67^. This suggests that articulatory control may rely on synergies or primitives, rather than the control of individual articulators or muscles^68^.

We have currently implemented the state estimation process in FACTS as an Unscented Kalman Filter. We intend this to be purely a mathematically tractable approximation of the actual neural computational process. Interestingly, recent work suggests that a related approach to nonlinear Bayesian estimation, the Neural Particle Filter, may provide a more neurobiologically plausible basis for the state estimation process^69^. Our future extensions of FACTS will involve exploring implementing this type of filter in FACTS.

In conclusion, the FACTS model uses a widely accepted domain-general approach to motor control, is able to replicate many important speech behaviors, and makes new predictions that can be experimentally tested. This model pushes forward our knowledge of the human speech motor control system, and we plan to further develop the model to incorporate other aspects of speech motor behavior, such as pitch control and sensorimotor learning, in future work.

## Methods

### Notation

We use the following mathematical notation to present the analyses described in this paper. Matrices are represented by bold uppercase letters (e.g., **X**), while vectors are represented in italics without any bold case (either upper or lower case). We use the notation **X**^*T*^ to denote the matrix transpose of **X**. Concatenations of vectors are represented using bold lowercase letters (e.g., **x** = [*x ẋ*]^*T*^). Scalar quantities are represented without bold and italics. Derivatives and estimates of vectors are represented with dot and tilde superscripts, respectively (i.e., **ẋ** and **x̃**, respectively).

### Task state feedback control law

In FACTS, we represent the state of the vocal tract tasks **x**_t_ = [*x*_*t*_ *ẋ*_*t*_]^*T*^ at time *t* by a set of constriction task variables *x*_*t*_ (given the current gestural implementation of speech tasks, this is a set of constriction degrees such as lip aperture, tongue tip constriction degree, velic aperture, etc. and constriction locations, such as tongue tip constriction location) and their velocities *ẋ*_*t*_. Given a gestural score generated using a linguistic gestural model^70, 71^, the *task state feedback control law* (equivalent to the Forward Task Dynamics model in^32^) allows us to generate the dynamical evolution of **x**_t_ using the following simple second-order critically-damped differential equation:

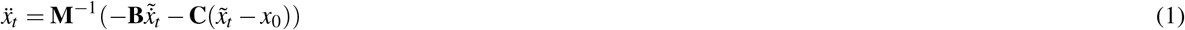

where *x*_0_ is the active task (or gestural) goal, **M**, **B**, and **C** are respectively the mass matrix, damping coefficient matrix, and stiffness coefficient matrix of the second-order dynamical system model. Essentially, the output of the task feedback controller, *ẍ*_*t*_, can be seen as a desired change (or command) in task space. This is passed to the articulatory state feedback control law to generate appropriate motor commands that will move the plant to achieve the desired task-level change.

Although the model does include a dynamical formulation of the evolution of speech tasks (following^32, 35^), this is not intended to model the dynamics of the vocal tract plant itself. Rather, the individual speech tasks are modelled as (abstract) dynamical systems.

### Articulatory state feedback control law

The desired task-level state change generated by the task feedback control law, *ẍ*_*t*_, is passed to an articulatory feedback control law. In our implementation of this control law, we use Eq. 2 (after^32^) to perform an inverse kinematics mapping from the task accelerations *ẍ*_*t*_ to the model articulator accelerations *ä*_*t*_, a process which is also dependent on the current estimate of the articulator positions *ã*_*t*_ and velocities *ȧ̃*_*t*_. **J**(*ã*) is the Jacobian matrix of the forward kinematics model relating changes in articulatory states to changes in task states, **J̇**(*ã*, *ȧ̃*) is the result of differentiating the elements of **J**(*ã*) with respect to time, and **J**(*ã*)* is a weighted Jacobian pseudoinverse of **J**(*ã*).
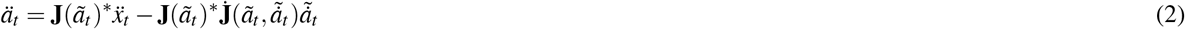

### Plant

In order to generate articulatory movements in CASY, we use Runge-Kutta integration to combine the previous articulatory state of the plant ([*a*_*t*−1_ *ȧ*_*t*−1_]^*T*^) with the output of the inverse kinematics computation (*ä*_*t*−1_, the input to the plant, which we refer to as the **motor command**). This allows us to compute the model articulator positions and velocities for the next time-step ([*a*_*t*_ *ȧ*_*t*_]^*T*^), which effectively “moves” the articulatory vocal tract model. Then, a tube-based *synthesis model* converts the model articulator and constriction task values into the output acoustics (parameterized by the vector 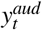). In order to model noise in the neural system, zero-mean Gaussian white noise *ɛ* is added to the motor command (*ä*_*t*−1_) received by the plant as well as to the somatosensory 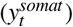 and auditory 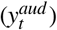 signals passed from the plant to the articulatory state estimator. Currently, noise levels (standard deviation of Gaussian noise) are tuned by hand for each of these signals (see below for details). Together, the CASY model and the acoustic synthesis process constitute the plant. The model vocal tract in the current implementation of the FACTS model is the Haskins Configurable Articulatory Synthesizer (or CASY)^38, 41, 42^.

### Articulatory state estimator

The articulatory state estimator generates an estimate of the articulatory state of the plant needed to generate state-dependent motor commands. The final state estimate 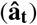 generated by the observer is a combination of an articulatory state prediction (**ã_t_**) generated from an efference copy of outgoing motor commands, combined with information about the state of the plant derived from the somatosensory and auditory systems (**y**_t_). This combination of internal prediction and sensory information is accomplished through the use of an Unscented Kalman Filter (UKF)^36^, which extends the linear Kalman Filter^28^ used in most non-speech motor control models^23, 25^ to nonlinear systems like the speech production system.

First, the state prediction is generated using a **forward model** 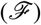 that predicts the evolution of the plant based on an estimate of the previous state of the plant (**ã**_t−1_) and an efference copy of the previously issued motor command (*ä*_*t*−1_). Based on this predicted state, another forward model (*ℋ*) generates the predicted sensory output 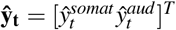 (comprising somatosensory and auditory signals 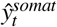 and 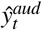, respectively) that would be generated by the plant in the predicted state. Currently, auditory signals are modelled as the first three formant values (F1-F3; 3 dimensions), and somatosensory signals are modelled as the position and velocities of the speech articulators in the CASY model (20 dimensions).

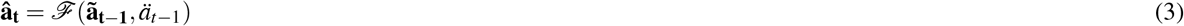

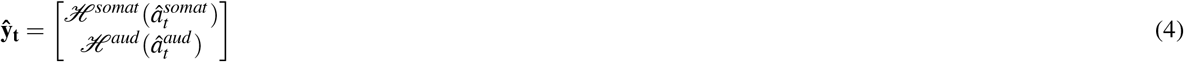

These predicted sensory signals are then compared with the incoming signals from the somatosensory 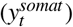 and auditory 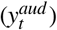 systems, generating the sensory prediction error (comprising both somatosensory and auditory components) 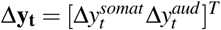:

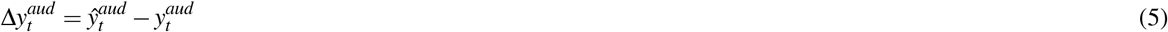

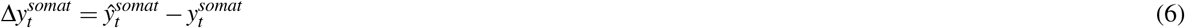

These sensory prediction errors are used to correct the initial articulatory state prediction, giving a final articulatory state estimate **ã**_t_:

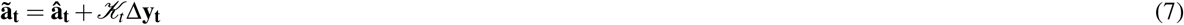

where 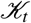 is the Kalman Gain, which effectively specifies the weights given to the sensory signals in informing the final state estimate. Details of how we generate 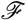, *ℋ*, and 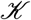 are given in the following sections.

### Forward models for state and sensory prediction

One of the challenges in estimating the state of the plant is that both the process model 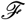 (that provides a functional mapping from [*ã*_*t*−1_ *ȧ̃*_*t*−1_ *ä*_*t*−1_]^*T*^ to **â**_t_) as well as the observation model *ℋ* (that maps from 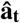 to ***ŷ***_t_) are unknown. Currently, we implement the process model 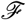 by replicating the integration of *ä*_*t*_ used to drive changes in the CASY model, which ignores any potential dynamical effects in the plant. However, the underlying architecture (the forward model 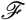) is sufficiently general that non-zero articulatory dynamics could be accounted for in predicting 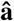 as well.

Implementing the observation model is more challenging due to the nonlinear relationship between articulator positions and formant values. In order to solve this problem, we *learn* the observation model functional mappings from articulatory positions to acoustics 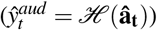) required for Unscented Kalman Filtering using Locally Weighted Projection Regression, or LWPR, a computationally efficient machine learning technique^37^. While we do not here explicitly relate this machine learning process to human learning, such maps could theoretically be learned during early speech acquisition, such as babbling^22^. Currently, we learn only the auditory prediction component of *ℋ*. Since the dimensions of the somatosensory prediction are identical to those of the predicted articulatory state, the former are generated from the latter via an identity function 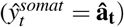.

### State correction using an Unscented Kalman Filter

Errors between predicted and afferent sensory signals are used to correct the initial efference-copy-based state prediction through the use of a Kalman gain (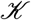, Eq. 7), which effectively dictates how each dimension of the sensory error signal Δ**y**_*t*_ affects each dimension of the state prediction **â**_t_. Our previous SFC model of vocal pitch control implemented a Kalman filter to estimate the weights in the Kalman gain^29^, which provides the optimal state estimate under certain strict conditions, including that the system being estimated is linear^28^. However, the traditional Kalman filter approach is only applicable to linear systems, and the speech production mechanism, even in the simplified CASY model used in FACTS, is highly non-linear.

Our goal in generating a state estimate is to combine the state prediction and sensory feedback to generate an optimal or near-optimal solution. To accomplish this, we use an Unscented Kalman Filter (UKF)^36^, which extends the linear Kalman Filter to non-linear systems. While the UKF has not been proven to provide optimal solutions to the state estimation problem, it consistently provides more accurate estimates than other non-linear extensions of the Kalman filter, such as the Extended Kalman Filter^36^.

In FACTS, the weights of the Kalman gain are computed as a function of the estimated covariation between the articulatory state and sensory signals, given by the posterior covariance matrices 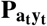 and 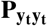 as follows:

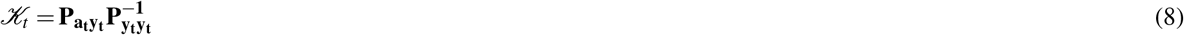

In order to generate the posterior means (the state and sensory predictions) and covariances (used to caculate 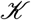) in an unscented Kalman Filter (UKF)^36^, multiple prior points (called sigma points, 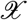) are used. These prior points are chosen carefully to capture the mean and covariance of the prior state. Each of these points is then projected through the nonlinear forward model function (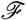 or *ℋ*), after which the posterior mean and covariance can be calculated from the transformed points. This process is called the unscented transform (Fig. 8). This is used both to predict the future state of the system (process model, 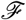) as well as the expected sensory feedback (observation model, *ℋ*).

**Figure 8.**
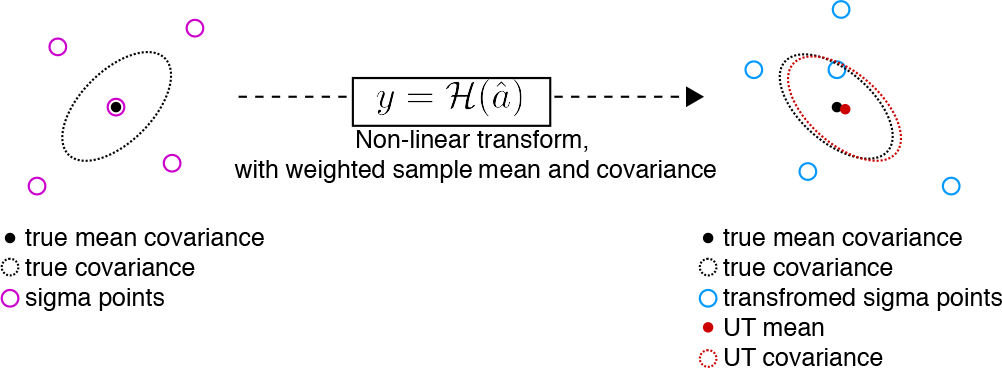
Cartoon showing the unscented transform (UT). The final estimate of the mean and covariance provide a better fit for the true values than would be achieved with the transformation of only a single point at the pre-transformation mean.

First, the sigma points 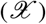 are generated:

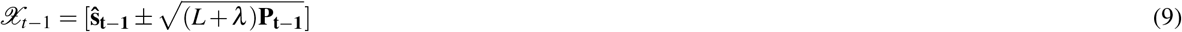

where 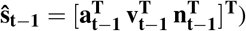 and **v** and **n** are estimates of the process noise (noise in the plant articulatory system) and observation noise (noise in the sensory systems), respectively, ***L*** is the dimension of the dimension of the articulatory state **a**, *λ* is a scaling factor, and **P** is the noise covariance of **a**, **v**, and **n**. In our current implementation, the level of noise for **v** and **n** are set to be equal to the level of Gaussian noise added to the plant and sensory systems.

The observer then estimates how the motor command *ä*_*t*_ would effect the speech articulators by replicating using the Euler integration model 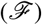 to generate the state prediction 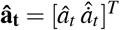. First, all sigma points reflecting the articulatory state 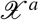 and process noise 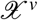 are passed through 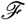:

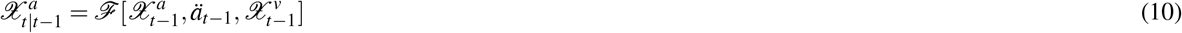

and the estimated articulatory state is calculated as the weighted sum of the sigma points where the weights (*W*) are inversely related to the distance of the sigma point from the center of the distribution.
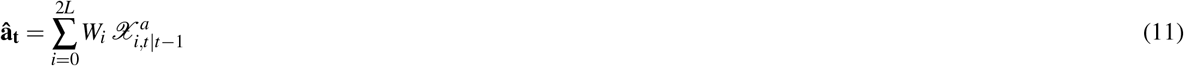

The expected sensory state (*ŷ*_*t*_) is then derived based on the predicted articulatory state in a similar manner, first by projecting the articulatory 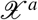 and observation noise 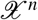 sigma points through the articulatory-to-sensory transform *ℋ*.
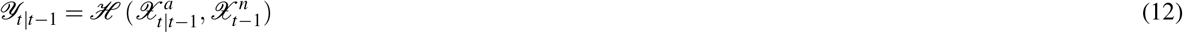

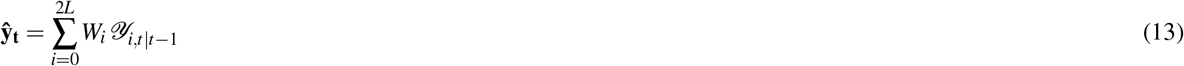

Lastly, the posterior covariance matrices 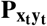 and 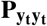 necessary to generate the Kalman Gain 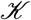 as well as the Kalman Gain itself are calculated in the following manner:

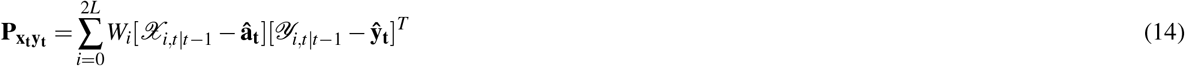

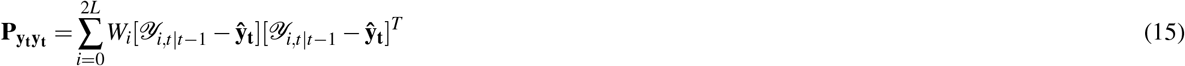

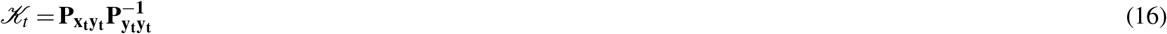

### Task state estimator

Finally, we estimate the vocal tract state estimate at the next time step by passing the articulatory state estimate into a task state estimator, which in our current implementation is a forward kinematics model (see Eq. 2)^32^. **J**(*ã*), the Jacobian matrix relating changes in articulatory states to changes in task states, is the same as in Eq. 2.
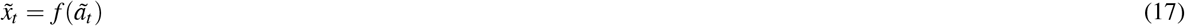

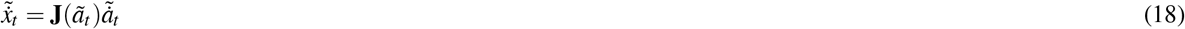

This task state estimate is then passed to the task feedback controller to generate the next task-level command *ẍ*_*t*_ using Eq.1.

### Model parameter settings

There are a number of tunable parameters in the FACTS model. These include: 1) the noise added to *ä* in the plant, *y*_*aud*_ in the auditory system, and *y*_*somat*_ in the somatosensory system; 2) the internal estimates of the process (*ä*) and observation **y** noise; and 3) initial values for the process, observation, and state covariance matrices used in the Unscented Kalman Filter. Internal estimates of the process and observation noise were set to be equal to the true noise levels. Noise levels were tuned by hand (using a range from 1e-1 to 1e-8, scaled by the norm of each signal) to achieve the following goals: stable system behavior in the absence of external perturbations, the ability of the model to react to external auditory and somatosensory perturbations, and a partial compensation for external auditory perturbations in line with observed human behavior. The final noise values used were 1e-4 for the plant/process noise, 1e-2 for the auditory noise, and 1e-6 for the somatosensory noise. The discrepancy in the values for the noise between the two sensory domains is proportional to the difference in magnitude between the two signals (300-3100 Hz for the auditory signal, 0-1.2 mm or mm/s for the articulatory position and velocity signals). Process and observation covariance matrices were initialized as identity matrices scaled by the process and observation noise, respectively. The state covariance matrix was initialized as an identity matrix scaled by 1e-2. A relatively wide range of noise values produced similar behavior: the effects of changing the auditory and somatosensory noise levels are discussed in the results section.

## Footnotes

1 Consistent with engineering control theory, we refer to the term “controller” as a “control law”.

